# Effects of co-incubation of LPS-stimulated RAW 264.7 macrophages on leptin production by 3T3-L1 adipocytes: A method for co-incubating distinct adipose tissue cell lines

**DOI:** 10.1101/2021.07.27.454005

**Authors:** Cristina Caldari-Torres, Jordan Beck

## Abstract

Adipose tissue is a major endocrine organ capable of releasing inflammatory adipokines. Inflammatory adipokine release is linked to the changes occurring in adipose tissue in the overfed state, where tissue remodeling results in hypertrophic adipocytes that recruit monocytes to infiltrate the tissue and take on an inflammatory phenotype. As the proportion of inflammatory macrophages increases there is a concurrent increase in release of macrophage-specific inflammatory mediators, further contributing to the inflamed state and setting an inflammatory loop between the macrophages and adipocytes. Although most inflammatory adipokines are released by macrophages, adipocytes can also release immunomodulatory adipokines, such as leptin. The objective of this research was to determine if co-incubation of activated macrophages with mature adipocytes, using Transwell inserts, affected leptin release by mature adipocytes. We also examined if there were differences in the amount of cell-secreted products quantified in cell-conditioned media collected from macrophage-containing (Transwell insert) and adipocyte-containing (well) compartments. Mature adipocytes (differentiated 3T3-L1 murine fibroblasts) were co-incubated with control and lipopolysaccharide-stimulated (0.01 μg/ml) murine macrophages (RAW264.7), and nitric oxide, interleukin-6, and leptin levels were quantified in the cell-conditioned media from the two compartments. Activation status of the macrophages did not affect leptin release by the adipocytes. We observed higher amounts of leptin in wells compared to Transwells. Nitric oxide and interleukin-6 levels were similar between Transwells and wells, suggesting that these adipokines are traveling through the Transwell inserts and reaching equilibrium between the two compartments. There was a weak negative relationship between nitric oxide release by macrophages and leptin release by adipocytes. Our results suggest that co-incubating activated macrophages and adipocytes using Transwell inserts can result in distinct microenvironments in the different cellular compartments and that separate sampling of these compartments is required to detect the subtle signaling dynamics that exist between these cells.

## Introduction

Adipose tissue (AT) has emerged as a major endocrine organ capable of releasing inflammatory mediators—adipokines—that may result in a chronic inflammatory state. Low levels of chronic inflammation, or meta-inflammation, may predispose the individual to chronic disease, including insulin resistance and type 2 diabetes mellitus (1,2). Adipokines are AT-derived proteins, a category that includes, but is not limited to, cytokines, chemokines, and hormones. In addition to adipocytes, AT contains adipocyte precursors in various stages of differentiation, as well as non-fat cells like endothelial and immune cells. Amongst the immune cells present in AT, macrophages have received a lot of attention by adipoimmunologists due to their central role in AT-derived inflammation.

Macrophages are innate immune cells that circulate as monocytes and differentiate into their final phenotype based on the microenvironment encountered in the tissue they extravasate into for residence. A comprehensive review of adipokine release from AT concluded that non-fat cells, mostly macrophages, release the majority of the inflammatory adipokines that are increased in the obese state (3). Some of the adipokines found at higher circulating levels in obese individuals include interleukin (IL)-6, IL-8, tumor necrosis factor (TNF)-α, monocyte chemoattractant protein (MCP)-1, and leptin (3). Of these adipokines, leptin is the only one that is primarily produced by the adipocytes (3). Leptin is positively correlated with obesity and has a major role in regulation of energy homeostasis (4) through its function as a signal in the feedback loop that controls food intake and body weight (5). Leptin also has effects on both the innate and adaptive branches of the immune system. With regards to monocytes/macrophages, which express the functional leptin receptor (6, 7), this adipokine has been implicated in modulation of cytokine production (7), phagocytosis (8), and differentiation of monocytes into pro-inflammatory M1 macrophages (9, 10). Unno et al. (11) reported that nitric oxide (NO), a signaling molecule that is readily produced by activated macrophages, downregulated leptin expression at both the protein and mRNA level in murine adipocytes (3T3-L1 cell line). The NO used in the study was derived from various NO donors (NOC7, NOC18, and GSNO) and autocrinally from the adipocytes. There are currently no reports on the effects of macrophage-derived adipokines, including NO, on leptin production by adipocytes.

The inflamed microenvironment caused by AT-derived adipokines in the obese state drives immune cell recruitment (3, 12, 13), increasing the percentage of macrophages residing in obese compared to lean AT. In lean AT, macrophages make up ~10% of cells (14) and mostly exhibit the alternatively activated (M2) anti-inflammatory phenotype. M2 macrophages have housekeeping functions ranging from immune surveillance to clearance of cellular debris and lipid buffering (15). In obese AT there is an increase in total macrophage numbers (making up to 50-60% of AT) as well as in the number of macrophages exhibiting a classically activated or M1, pro-inflammatory phenotype (14, 16). Pro-inflammatory M1 macrophages express higher levels of TNF-α and inducible nitric oxide synthase (iNOS) (16). Using DNA microarray gene analyses, Yamashita et al. (17) concluded that low levels (1 ng/ml) of bacterial lipopolysaccharide (LPS) drive RAW264.7 murine macrophages to differentiate into M1 macrophages, increasing expression of cyclooxygenase-2, iNOS, TNF-α, and activation of NF-κB. Factors that increase monocyte recruitment to obese AT and their differentiation into the M1 macrophage phenotype include hypertrophy of adipocytes (18), the aforementioned release of inflammatory adipokines and chemokines by AT (19, 20), and increased presence of hypoxic/apoptotic adipocytes (21). Adipocyte survival and maturation/differentiation is affected by increases in M1 macrophage numbers in AT, resulting in a decrease in adipocyte hyperplasia--the production of new adipocytes--during times of chronic positive energy balance, and instead favoring hypertrophy of already existing adipocytes (2). Hypertrophic adipocytes are associated with augmented inflammation and dysfunctional insulin sensitivity (22). Inflammatory adipokines released by M1 macrophages, like TNF-α and IL-6, can block insulin action in adipocytes via autocrine/paracrine mechanisms (12), linking the increased macrophage recruitment and M1 polarization observed in obese AT with impaired insulin sensitivity. These inflammatory adipokines also result in adipocyte mitochondrial dysregulation, through an increased release of reactive oxygen species and mitochondrial fragmentation, adding a further layer of complexity to the adipocyte-macrophage crosstalk and potentiation of inflammation in obese adipose tissue (23).

In review, increased energy storage associated with obesity causes hypertrophic, hypoxic, and apoptotic adipocytes that release increasing amounts of inflammatory adipokines. The inflamed microenvironment favors recruitment of macrophages toward obese AT and polarizes them toward the M1 inflammatory phenotype which, in turn, release macrophage-specific inflammatory adipokines that further support adipocyte hypertrophy and recruitment of monocytes from circulation, creating an inflammatory loop. The cross-talk between macrophages and adipocytes and their precursors is central to the investigation of AT-derived inflammation, as it maintains the inflammatory loop and aids in the recruitment of new macrophages that will likely develop an M1 phenotype (13).

The main objective of this research was to determine the effects of co-incubating murine 3T3-L1 adipocytes and activated RAW264.7 macrophages on the production of two inflammatory adipokines--IL-6 and leptin—by these two cells types. Specifically, we wanted to test if activation status of the macrophages would exert paracrine effects on the mature adipocytes, as measured by secretion of leptin, an adipocyte-specific adipokine. These objectives were tested through the use of Transwell inserts (0.4 μm pore size), which allow for the co-incubation of different cell lines and exposure of one cell line to products secreted by the other cell line. We also examined if there was a difference in the amount of cell-secreted products quantified in the cell-conditioned media collected from macrophage-containing Transwells and adipocyte-containing wells. Sampling each cell types’ microenvironment would allow us to detect the potential subtle signaling dynamics that exist between these cells.

## Materials and Methods

### Reagents and materials

Murine fibroblast (3T3-L1, cat no. CL-173) and macrophage (RAW 264.7, cat no. TIB-71) immortalized cell lines were purchased from ATCC (Manassass, VA). Dulbecco’s modified Eagle’s medium (DMEM), phosphate buffered saline (PBS), fetal bovine serum (FBS), penicillin-streptomycin, and polystyrene 6-well plates were purchased from Fisher Scientific (Pittsburg, PA). The IL-6 and leptin enzyme-linked immunosorbent assay (ELISA) was purchased from R&D systems (Minneapolis, MN), while the Griess assay for nitric oxide (NO) quantification was obtained from Promega (Madison, WI). Trypan blue, insulin, dexamethasone (DEX), d-biotin, 3-isobutyl-1-methylxanthine (IBMX), Trypsin-EDTA 0.25%, and lipopolysaccharide (LPS) were purchased from Sigma-Aldrich (St Louis, MO). Transwell permeable supports (0.4 μm pore size, 12mm diameter, polyester membrane) and 12-well plates (polystyrene) were obtained from Corning Costar (Corning, NY).

### 3T3-L1 cell maintenance, culture, and differentiation into mature adipocytes

The 3T3-L1 cell line must be differentiated from a fibroblast phenotype into its final, mature adipocyte phenotype containing lipid droplets. To do this, cells were incubated in 12-well plates, in a 5% CO_2_ humidified atmosphere, and were kept in the undifferentiated fibroblast phenotype at less than 50%confluency during sub-culturing. Detachment of cell monolayer for sub-culturing was performed via trypsinization. Growth medium for 3T3-L1 cells consisted of DMEM, 10% (v/v) heat-inactivated FBS, 1% antibiotics (100 U/ml penicillin and 100 μg/ml streptomycin), and 0.008 μg/ml D-biotin. Differentiation into the adipocyte phenotype was performed as described by Zebisch *et al.* (24). Briefly, three days after cells reached ~90%confluency and started to clump together and lose fibroblast morphology, they were washed with 1X PBS and treated with a differentiation cocktail consisting of growth medium supplemented with 0.05 μM IBMX, 1 μM DEX, and 20 μg/ml insulin. Forty-eight hours after addition of the differentiation cocktail, the cells were washed with 1X PBS and treated with post-differentiation medium consisting of growth medium supplemented with 20 μg/ml insulin. Treatment with post-differentiation medium was performed every forty-eight hours for a total of four times. At the end of the differentiation period lipid droplets inside the adipocytes could be visualized using an inverted microscope.

### RAW 264.7 cell maintenance and culture

RAW 264.7 cells were grown in polystyrene 6-well plates with DMEM supplemented with 10% (v/v) heat-inactivated FBS and 1% antibiotics (100 U/ml penicillin and 100 μg/ml streptomycin). Cells were incubated in a 95% O_2_ and 5% CO_2_ humidified atmosphere. During initial expansion, the medium was changed every two days after washing cells with 1X PBS. Cells were not grown beyond 80%confluency during expansion. Detachment of cell monolayer for sub-culturing was performed with the cell scraping method. When cells reached 80%confluency, they were transferred to Transwell inserts to commence the co-incubation experiments.

### RAW 264.7 cell activation

A concentration of 0.01 μg/ml of LPS was used and three methods of LPS challenge for co-incubation experiments were tested: macrophages in 6-well plate were washed, resuspended (via scraping) in fresh medium, transferred to Transwell inserts and challenged with LPS that was added into the Transwell compartment (3T3+RAW+LPS); macrophages were challenged with LPS for 24 h while in well of 6-well plate, were then washed, resuspended (via scraping) in fresh medium, and transferred to Transwell insert (3T3+StimRAW); and macrophages were challenged with LPS for 24 h while in well of 6-well plate and then resuspended (via scraping) and transferred to Transwell insert along with the conditioned media (3T3+StimRAW+CondMed). Non-LPS-challenged macrophages were co-incubated with adipocytes as a control (3T3+RAW).

### Quantification of Nitric Oxide (NO) Production

Macrophage activation was quantified via measurement of NO levels. Nitric oxide production by the macrophages was determined through quantifying nitrite levels in cell-conditioned media using the Griess assay. Briefly, 50 μL of cell conditioned media were added in triplicate to a 96-well plate and mixed with 50 μL of sulfanilamide solution and allowed to incubate for 10 min. Following the incubation, 50 μL of N-1-naphthylethylenediamine (NED) were added to each well, followed by a 10 min incubation. After the second incubation, absorbance was measured at 530 nm. Nitrite concentrations were determined by extrapolating absorbance measurements from a 0-100 μM standard curve. An Epoch plate reader (BioTek Instruments, Winooski, VT) was used for absorbance measurements.

### Co-incubation of RAW 264.7 and 3T3-L1 cells using Transwell Inserts

Several co-incubation methodologies were tested to determine if they resulted in different activation level of the macrophages, as measured by NO and IL-6 levels in cell-conditioned medium. Additionally, leptin levels in cell-conditioned medium were measured to test if activation state of the macrophages affected production of this adipokine by the mature adipocytes. Three methods of macrophage stimulation/co-incubation were tested. Mature adipocytes (differentiated according to the steps described in section 2.2) in 12-well plates were co-incubated with macrophages stimulated with LPS as described in the section 2.4: 1. 3T3+RAW+LPS, 2. 3T3+StimRAW, and 3. 3T3+StimRAW+CondMed. Additionally, adipocytes were co-incubated with unstimulated RAW264.7 cells resuspended in fresh medium (3T3+RAW) as a control. On average, 8.0×10^5^ macrophages were plated onto each Transwell insert in a total of 500 μl. Twenty-four hours after co-incubation commenced, media was collected separately from the Transwell inserts and wells, transferred to 1.5 ml microcentrifuge tubes, and stored at −20°C until used for IL-6, leptin, and NO quantification. Wells were run in duplicate and experiments were performed four times.

### Quantification of Adipokine Production

Analyses of cell conditioned media for determination of adipokine levels were done using IL-6 and leptin sandwich ELISAs according to manufacturer’s instructions. Samples were tested in triplicate, and a standard curve was produced and used to extrapolate the cytokine concentrations in the samples. Samples measuring > 500 pg/ml (highest standard) were diluted, re-quantified, and results were adjusted taking the dilution factor into account. An Epoch plate reader was used for absorbance measurements.

### Statistical analyses

Statistical analyses were done using JMP Pro 13 (Cary, NC). Non-normal data were normalized using a log transformation. Matched paired t-tests were used to determine differences in NO, IL-6, and leptin levels between cell-conditioned medium collected from Transwell inserts containing macrophages and wells containing adipocytes. In order to determine differences in NO, IL-6, and leptin produced by control and LPS-challenged cells, Student’s t-tests were used. Linear regression analyses were used to test relationships between IL-6, NO, and leptin. Interleukin-6, leptin, and NO concentrations in cell conditioned media were analyzed using the general mixed linear model. All statistical analyses were conducted using JMP Pro 15 (SAS, Cary, North Carolina). The sources of variation included experiment, treatment, experiment x treatment interaction, and well nested within experiment x treatment interaction. The experiment, treatment x experiment interaction, and well nested within experiment x treatment interaction were considered as random variables. When treatment effects were detected, means were separated using Tukey’s HSD. The level of significance was defined at *p* < 0.05. Experimental results are expressed as mean ± SE.

## Results

### Differential effects of LPS on NO, IL-6, and leptin production in murine macrophage and adipocytes

In order to determine the effect of an LPS challenge on macrophage activation (as measured by NO production), IL-6, and leptin production by murine macrophages and adipocytes, a series of control experiments were performed on the isolated cell lines. The RAW 264.7 macrophages were grown to 80-85% confluence, while 3T3-L1 cells were differentiated into the mature adipocyte phenotype, at which time LPS was added (0.01 μg/ml) and allowed to incubate for a period of 6 hours. Lipopolysaccharide challenge resulted in an increase in macrophage activation, as measured by increased NO production (Figure 1a; student’s t-test, p < 0.0001), and IL-6 production (Figure 1b; student’s t-test, p < 0.0001), and increased IL-6 production by mature adipocytes (Figure 1b; t-test, p=0.001). Nitric oxide production by mature adipocytes was negligible and did not differ between control and LPS-challenged cells (Figure 1a; student’s t-test, p = 0.58), while murine macrophages did not produce quantifiable amounts of leptin in either the control or LPS-challenged conditions (Figure 1c). Adipocytes produced similar amounts of leptin in either the absence or presence of LPS (965.8 ± 648.2 vs 741.5 ± 413.2 pg/ml) (Figure 1c; student’s t-test, p = 0.43).

**Fig 1.**
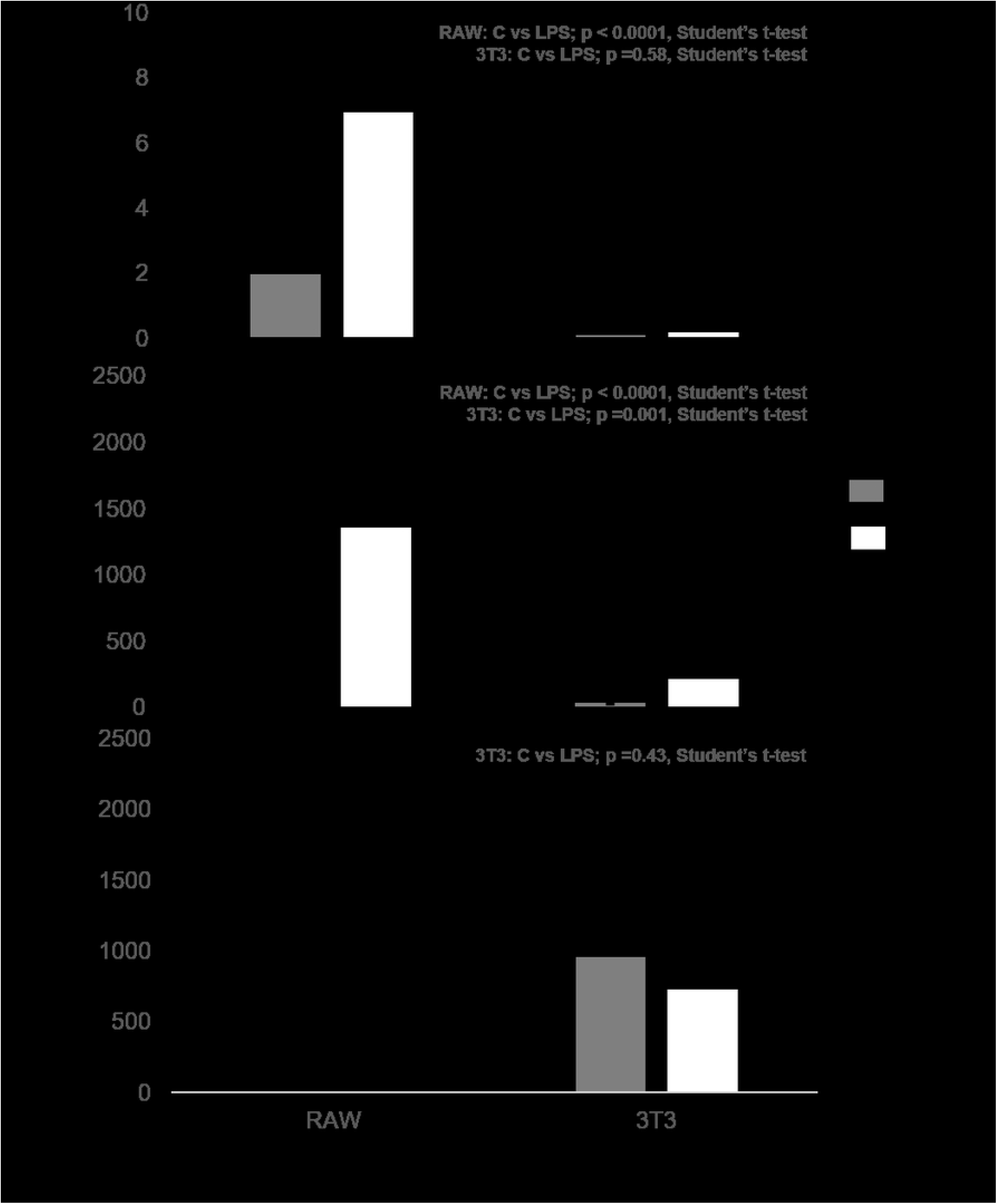
Nitric oxide (NO; μM, panel a), interleukin-6 (IL-6; pg/ml, panel b), and leptin (pg/ml, panel c) production by control and LPS-stimulated murine macrophages (RAW264.7) and adipocytes (3T3-L1). Adipokine concentrations in control and LPS-stimulated cells were compared using a Student’s t-test. Data represent least squares means ± SEM of 4 independent experiments. There was no quantifiable leptin production by RAW 264.7 cells. Significant treatment differences are represented with an asterisk (*) (p<0.05).

### Differences in amounts of molecules quantified between Transwell inserts and wells

Although the pores of the Transwell insert membrane are large enough (0.4 μm) to allow the diffusion of molecules of varying sizes from areas of higher to lower concentration, we considered that at the time of sampling equilibrium might not have been reached, resulting in differences in microenvironment between the Transwell inserts and the wells. At the end of the co-incubation experiments we sampled cell-conditioned media from these two compartments and measured NO, IL-6, and leptin levels in them separately (Table 1). Higher amounts of the leptina dipokine were observed in adipocyte-containing wells (627.1 ± 47.5 pg/ml) compared to macrophage-containing Transwell inserts (478.2 ± 47.5 pg/ml) (Table 1; matched pairs t-test, p = 0.004). These findings are unsurprising as leptin is mostly, if not exclusively, produced by adipocytes, which were localized in the well compartment. The difference in leptin levels in wells and Transwells indicates that if leptin is traveling across the Transwell insert membrane, at the time of sampling levels of the molecule had not reached equilibrium. There were no significant differences in the amounts of NO and IL-6 quantified in the macrophage-containing Transwell inserts compared to the adipocyte-containing wells (Table 1). The low levels of NO and IL-6 produced by control and LPS-challenged adipocytes (Figures 1a and 1b) suggest that these molecules are produced by macrophages and traveling across the Transwell membrane into the well compartment.

**Table 1.** Differences in NO, IL-6, and leptin amounts quantified between macrophage-containing Transwell inserts and adipocyte-containing wells (across treatments). Matched Pairs t-tests with p<0.05 indicate significant difference in amount of molecule measured in Transwells and wells.

| molecule | Average amount of molecule in Transwell (Macrophages) | Average amount of molecule in Well (Adipocytes) | Difference between Transwell and Well (Matched Pairs t-test) |
| --- | --- | --- | --- |
| NO ($\mu\text{M}$) | $7.2 \pm 1.0$ | $6.0 \pm 1.0$ | $p=0.26$ |
| IL-6 (pg/ml) | $1172.2 \pm 122.0$ | $1115.76 \pm 122.0$ | $p=0.65$ |
| Leptin (pg/ml) | $478.2 \pm 47.5$ | $627.1 \pm 47.5$ | $p=0.004^*$ |

### Effect of macrophage activation status on NO, IL-6, and leptin levels

We were interested in testing if different methods of macrophages activation resulted in quantifiable differences in NO and IL-6 production. Specifically, we challenged macrophages with LPS (6 h) at time of co-incubation with adipocytes (3T3+RAW+LPS) or challenged them with LPS for 6 h before co-incubation with adipocytes and then either co-incubated the adipocytes with the previously LPS-challenged macrophages resuspended in fresh media (3T3+StimRAW) or in their conditioned media (3T3+stimRAW+CondMed).

As expected, the co-incubation methodologies with LPS-challenged macrophages (3T3+RAW+LPS,) resulted in higher NO (Figures 2b) and IL-6 (Figures 3b and 3c) production compared to the control with unstimulated macrophages (3T3+RAW) (general mixed linear model, p<0.0001). When measuring adipokine levels in cell-conditioned media from Transwell inserts, we observed no differences in levels of molecules that indicate macrophage activation (NO, IL-6) among the three co-incubation methodologies containing LPS-challenged macrophages, regardless of if the cells were activated before or during plating onto the Transwell inserts, or if fresh or conditioned media was used (Figures 2b, 2c, 3b and 3c; general mixed linear model, p<0.001). Interestingly, in the well compartment there were no differences in NO levels among control and the LPS-challenged macrophage treatments (Figure 2c; general mixed linear model, p=0.05), while IL-6 levels were significantly higher in those treatments containing activated macrophages (Figure 3c; general mixed linear model, p<0.0001).

**Fig 2.**
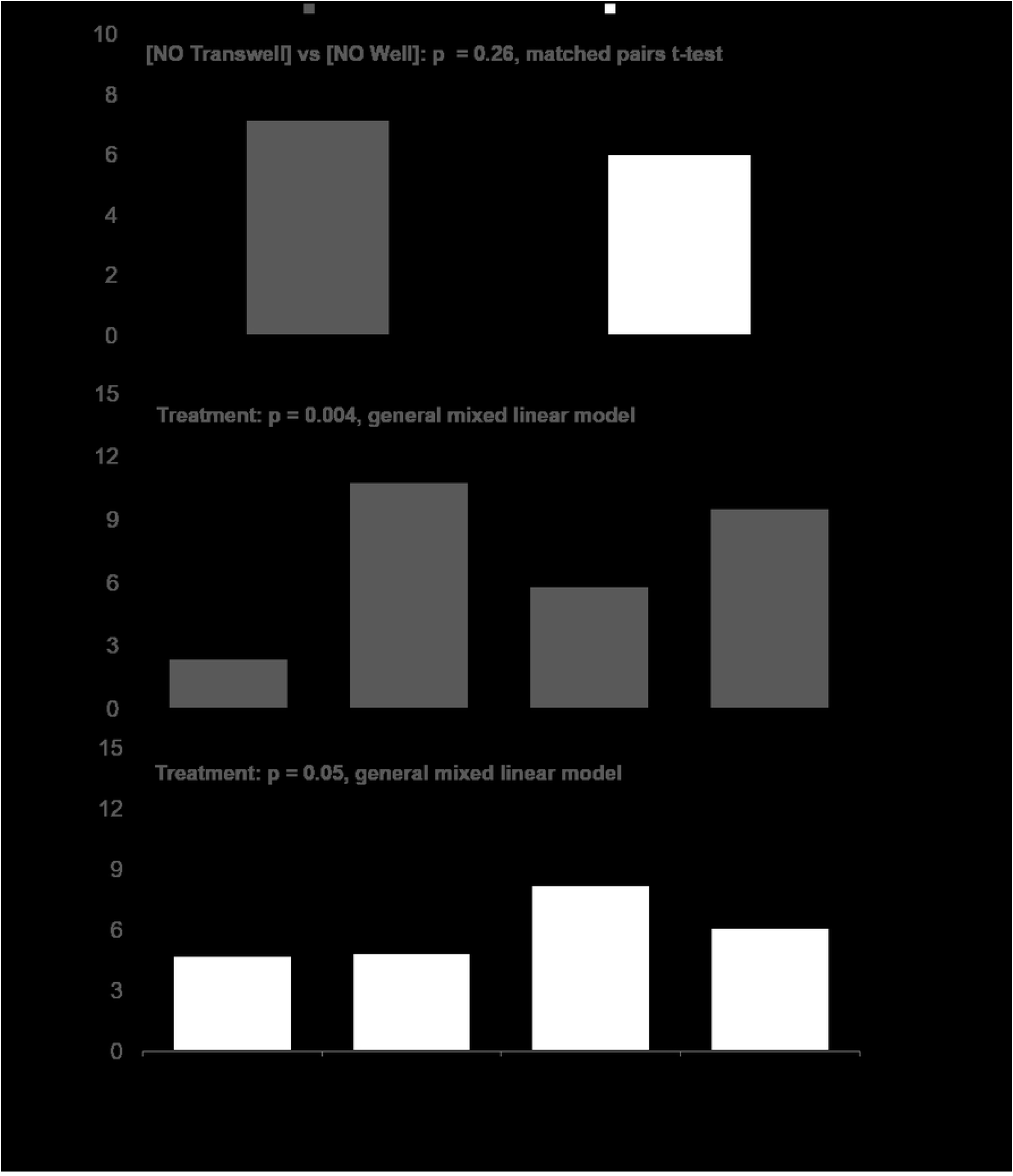
Mean nitric oxide (NO; μM) production across treatments in Transwells and wells (panel a) and for different co-incubation methods of 3T3-L1+RAW cells (Transwells=panel b; wells=panel c). NO levels in macrophage-containing Transwell inserts vs adipocyte-containing wells did not differ (matched-pairs t-test, p=0.26; panel a). NO concentrations for the various co-incubation methods (“treatment”) were compared using a general mixed linear model, followed by post-hoc Tukey-Kramer HSD. Data represent least squares means ± SEM of 4 independent experiments. Significant treatment differences are represented with different letters (p<0.05).

**Fig 3.**
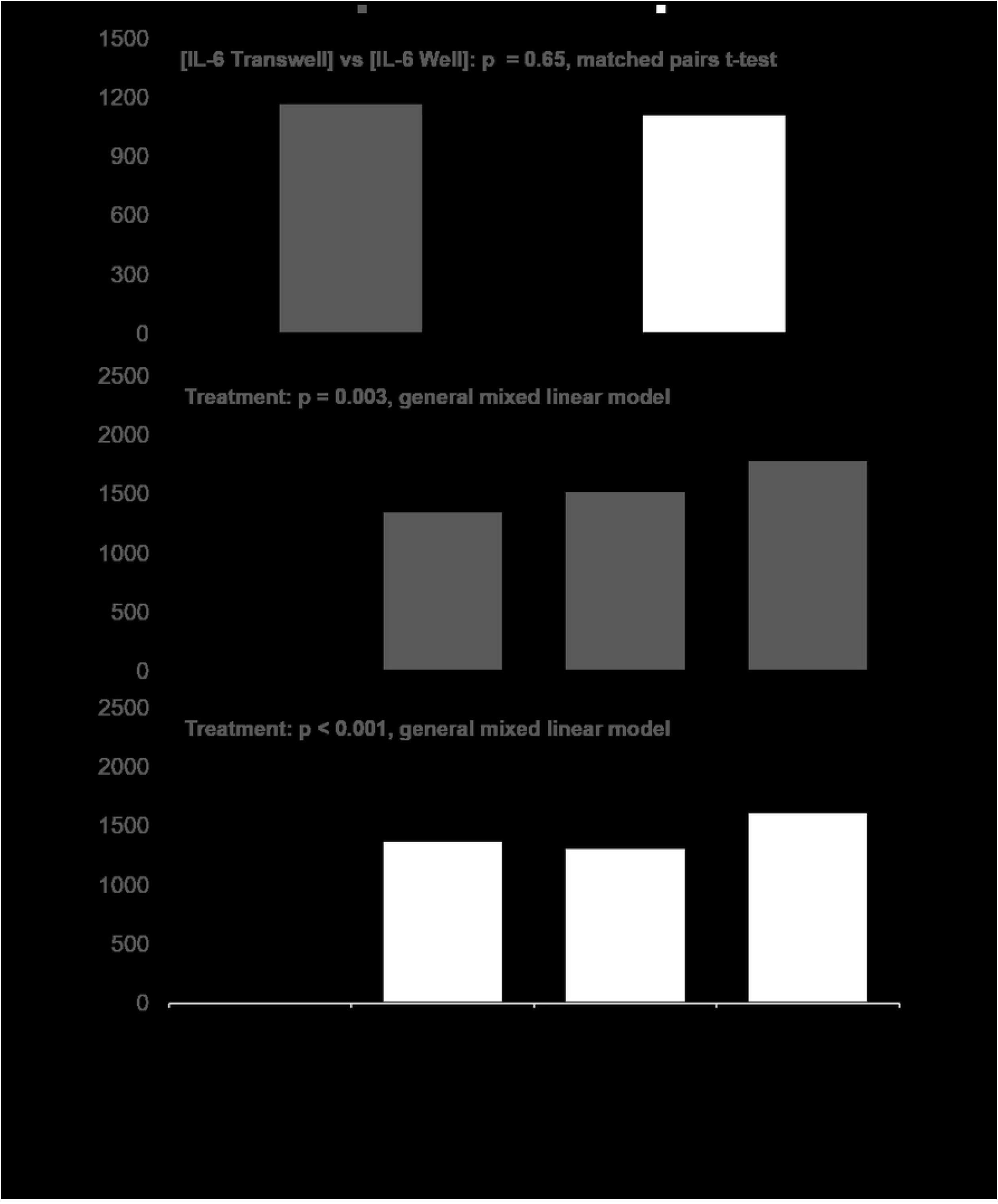
Mean interleukin-6 (IL-6; pg/ml) production across treatments in Transwells and wells (panel a) and for different co-incubation methods of 3T3-L1+RAW cells (Transwells=panel b; wells=panel c). IL-6 levels in macrophage-containing Transwell inserts vs adipocyte-containing wells did not differ (matched-pairs t-test, p=0.65; panel a). IL-6 concentrations for the various co-incubation methods (“treatment”) were compared using a general mixed linear model, followed by post-hoc Tukey-Kramer HSD. Data represent least squares means ± SEM of 4 independent experiments. Significant treatment differences are represented with different letters (p<0.05).

Higher amounts of the leptin adipokine were observed in adipocyte-containing wells compared to macrophage-containing Transwells (Fig 1a) but macrophage activation status did not have an effect on leptin levels in either the macrophage-containing Transwell inserts (Figure 4b) or adipocyte-containing wells (Figure 4c).

**Fig 4.**
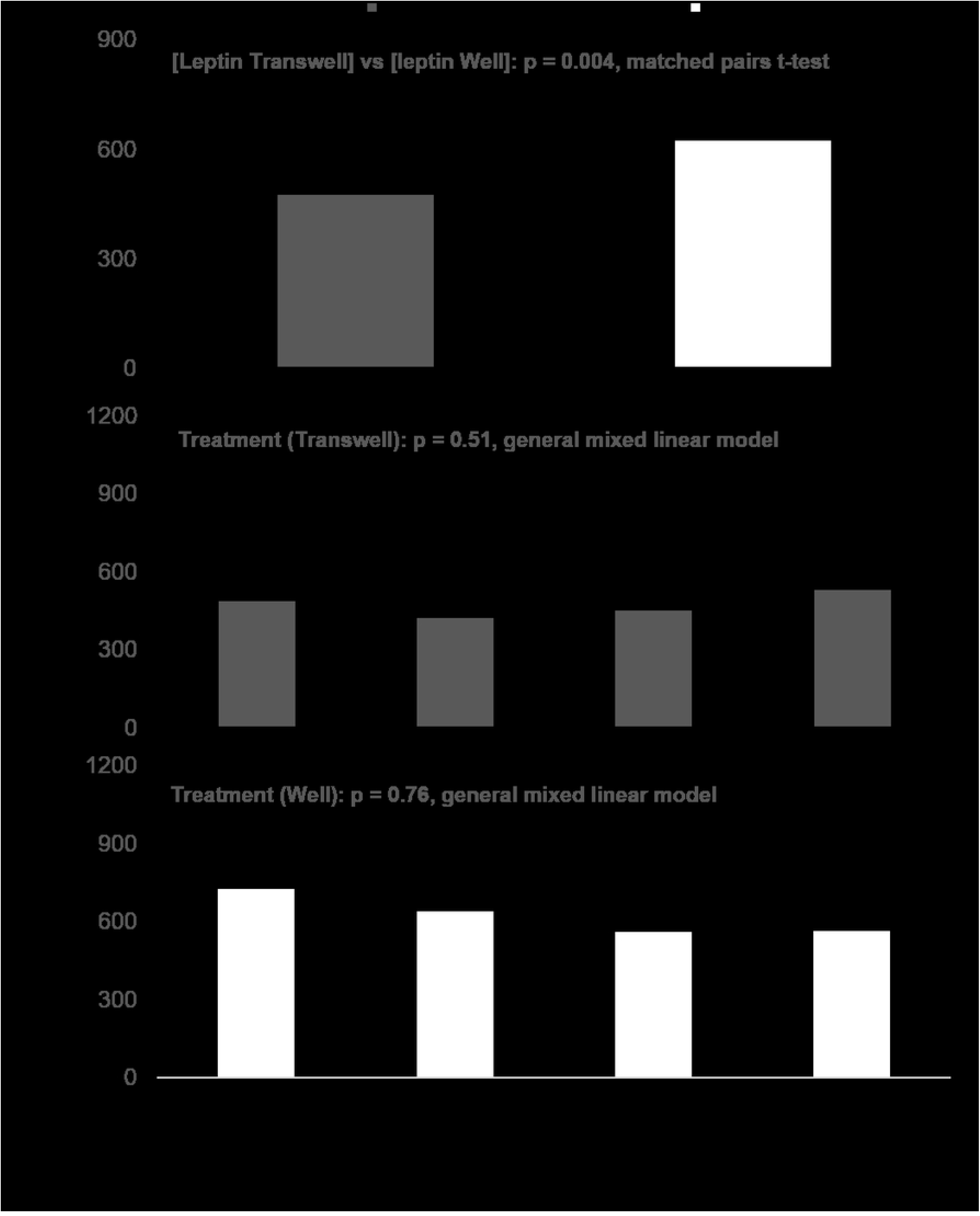
Mean leptin (pg/ml) production in Transwells and wells (panel a) and for different co-incubation methods of 3T3-L1+RAW cells (panels b and c). Leptin levels in macrophage-containing Transwell inserts were lower than for adipocyte-containing wells (matched-pairs t-test, p=0.004; panel a). Leptin concentrations for the various co-incubation methods (“treatment”) were compared using a general mixed linear model, followed by post-hoc Tukey-Kramer HSD. Data represent least squares means ± SEM of 4 independent experiments. There were no treatment effects on leptin levels in Transwell inserts or wells.

### Relationships between molecules

In Transwell inserts a trend for a negative linear relationship between levels of NO and leptin was observed (Figure 5a; linear regression, p=0.09), while a significant negative linear relationship between levels of NO and leptin was present in wells (Figure 5c; linear regression, p=0.03). There was no relationship between leptin and IL-6 levels in either wells or Transwell inserts (Figures 5b and 5d). It is important to highlight that the low coefficients of determinations (R^2^) for the relationships between NO and leptin indicate that there are high levels of variability in leptin levels that cannot be explained by NO levels.

**Fig 5.**
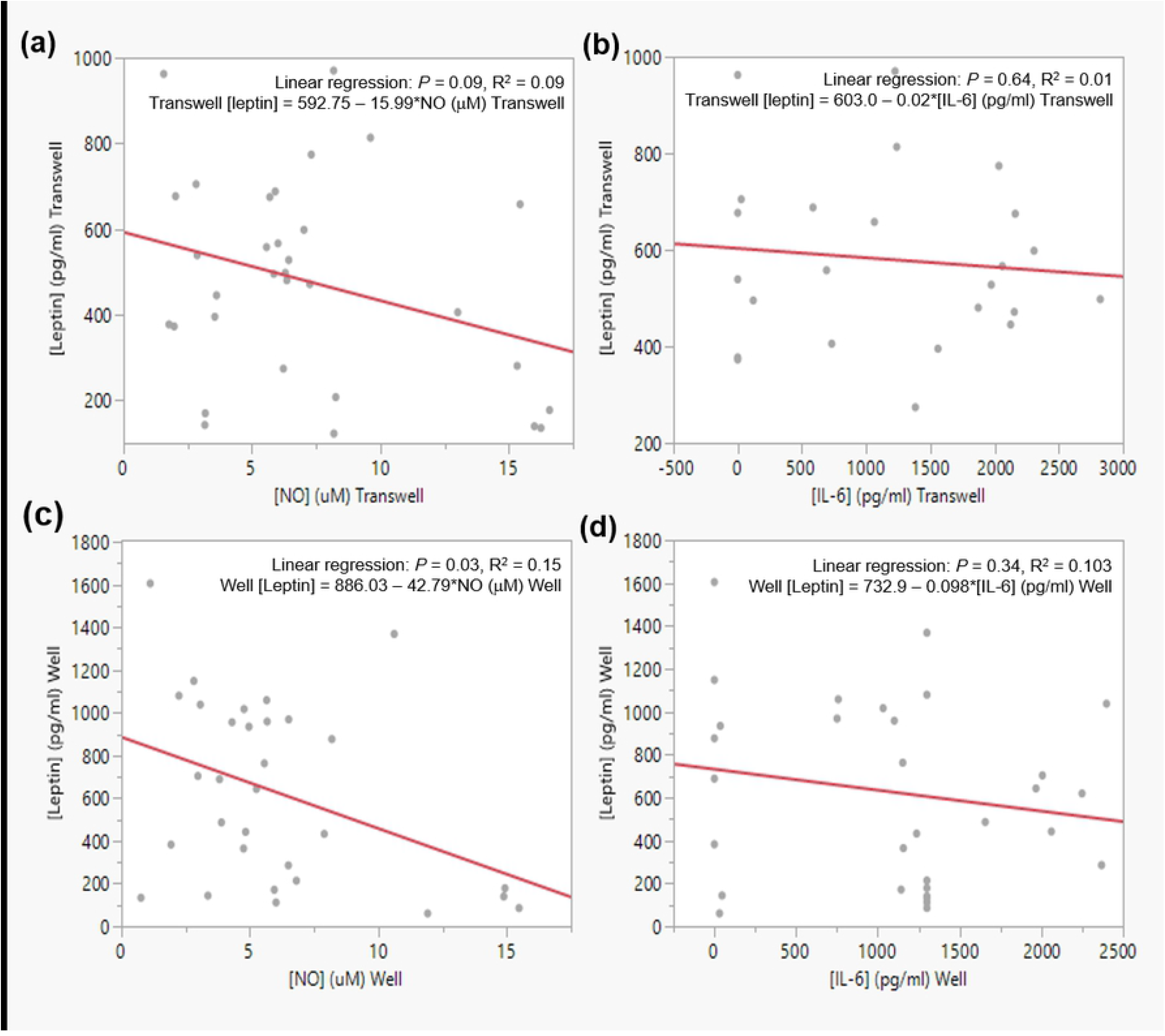
Relationship between NO and leptin and IL-6 and leptin concentrations measured in Transwell inserts (a and b) and wells (c and d). We observed a trend for a negative relationship between NO and leptin levels measured in Transwells (panel a; p=0.09, linear regression), and a significant negative relationship between these two molecules in wells (panel c; p=0.03, linear regression). There was no significant relationship between IL-6 and leptin levels in either Transwells (panel b) or wells (panel d).

## Discussion

There were significant differences in amounts of NO and leptin measured in Transwells and wells. Higher NO levels were observed in cell-conditioned media collected from the macrophage-containing Transwells, while the cell-conditioned media collected from adipocyte-containing wells contained higher levels of leptin. This is logical, as macrophages are the major contributors to NO while adipocytes are the major contributors to leptin in the cell-conditioned media. The assay used to determine NO levels measures concentrations of nitrite (NO^−^_2_), a small molecule (46 Da) that would be expected to travel through the pores of the Transwell membrane (0.4 μm size). Diffusion of nitrite through the pores of the inserts would explain why there were quantifiable amounts of this molecule in the adipocyte-containing wells. Unno et al. (11) reported that 3T3-L1 adipocytes treated with an interferon (IFN)-γ-LPS (10 ng/ml-5 μg/ml) mixture for 24 h had significantly increased expression of inducible nitric oxide synthase (iNOS) as well as of nitrite and nitrate released into culture medium, compared to untreated adipocytes. Dobashi et al. (25) treated differentiated 3T3-L1 adipocytes with 1 μg/ml LPS for 48 h and saw no significant increase in NO production by adipocytes compared to non-LPS treated cells, suggesting that IFN-γdrives iNOS expression in 3T3-L1 cells to a larger extent than LPS. Our results support this idea, as in control experiments we did not observe a significant effect of LPS challenge on adipocyte-derived NO production (Figure 1b). Taking these results into account we hypothesize that in our co-incubation system the presence of NO in wells is due to diffusion of macrophage-derived NO from the Transwell inserts into the wells.

Despite the differences in NO measured between Transwells and wells, IL-6 levels in cell-conditioned media collected from these two compartments were similar. Adipocytes produce IL-6 under the action of LPS (26) but it is unlikely that the lack of difference in IL-6 levels between the Transwells and wells is due to adipocyte-secreted IL-6, since the control experiments showed that LPS-challenged adipocytes produced about a sixth the amount of IL-6 under the action of LPS compared to macrophages (RAW: 1371.8 ± 682.9; 3T3: 225.9 ± 73.4 pg/ml; Figure 1b). We also suspect that LPS is not able to travel across the membrane of the Transwell insert, where it was added to the macrophages. Although the small size of the LPS molecule (4.3 kDa) means that it is possible for it to travel across the Transwell insert membrane, its heterogeneous nature can result in aggregates of varying sizes. These aggregates can range in size from 1000-4000 kDa (27), which would be too large to travel through the 0.4 μm pore size of the membrane. It is more plausible that IL-6, with a 21 kDa size, is small enough to diffuse across the Transwell membrane which would allow for equilibrium to be reached between the macrophage and adipocyte-containing compartments. The possibility that IL-6 production by macrophages could start leveling off before that of NO would explain the difference in NO levels observed between Transwells and wells, and the lack of difference observed for IL-6. We should consider that if LPS is traveling across the Transwell membrane it could be having a more pronounced effect on the co-incubated adipocytes than demonstrated in the control experiments that tested the effects of this endotoxin on the isolated cell lines. Yamashita et al. (28) found that IL-6 production was markedly up-regulated in adipocytes co-cultured with macrophages in the presence of LPS, compared to stimulating each cell line separately with the endotoxin. It is unlikely that in our system the IL-6 quantified in the wells is derived from the adipocytes, as levels of this adipokine are similar in cell-conditioned media from wells containing adipocytes co-incubated with macrophages in the presence of LPS (3T3+RAW+LPS) and wells containing adipocytes co-incubated with previously LPS-activated macrophages that were washed and resuspended in fresh medium (removing LPS) at the time of co-incubation (3T3+StimRAW).

The adipocyte-specific adipokine leptin was found in higher quantities in the wells, where the adipocytes resided. Temporal differences in expression of the leptin gene and production of the protein could explain the difference in Transwell vs well concentrations. The leptin molecule, at 16 kDa, is smaller than IL-6, which would allow it to travel across the Transwell membrane. If leptin secretion by the adipocytes is occurring 24 h post co-incubation, this would explain why levels of this adipokine are different between Transwells and wells during collection of cell-conditioned media. Our data show that there is basal leptin production by the adipocytes that is not dependent on LPS stimulation, as adipocytes exposed to the control treatment secreted the same amount of this adipokine as adipocytes co-incubated with LPS. Previous reports have found that both physiological and pathological levels of leptin do not induce the expression of IL-6 in murine macrophages, but that it augments the effect of LPS in inducing IL-6 expression by priming macrophages to be more responsive to this endotoxin and that this synergistic effect is mediated by interleukin-receptor associated kinase (IRAK)-1 (29). It is difficult to extrapolate these results to our own, since in our system we did not observe a difference in IL-6 levels between control experiments (macrophages activated with LPS) and macrophages in the co-incubation system which were activated with LPS and exposed to adipocyte-derived leptin. Furthermore, activation status of the macrophages did not affect leptin production although there appeared to be a weak relationship between leptin and NO levels, with leptin levels decreasing as NO levels increased. These results match what was observed by Unno et al. (11), who treated differentiated 3T3-L1 cells with an INF-γ-LPS mixture and observed a significant induction of iNOS and decrease in leptin at both the protein and mRNA levels. We need to take into account that despite the observed significant relationship between these molecules, the low R^2^ values indicate that there are other factors influencing this relationship. Use of INF-γ, in addition to LPS, in these co-incubation experiments could help clarify the relationship between leptin and NO in adipocytes. We also need to consider that the NO present in our system is mostly derived from macrophages, while in Unno’s system NO was produced autocrinally by the adipocytes or due to the synergistic effect of LPS and INF-γ. Nitric oxide might have differential autocrine and paracrine effects on leptin protein and gene expression in adipocytes. This is a question that warrants further investigation in order to better understand the effects macrophage-derived NO could potentially have on mature adipocytes in this co-incubation system.

As a methodology, co-incubation of activated macrophages and fully differentiated adipocytes can help answer questions about macrophage-adipocyte interactions in AT and provide insights into how to blunt the inflammatory loop observed in obese AT. Research on this subject suggests that monocyte/macrophage recruitment into obese AT is an early contributor to this loop by virtue of macrophages having a bigger role in the secretion of inflammatory adipokines compared to adipocytes (3, 13). The polarization of macrophages toward the M1 phenotype as they arrive in obese AT is presumably directed by the microenvironment encountered by the macrophages, which is mostly set by the adipocytes residing in the tissue (3, 30, 12). We wanted to examine if activation of macrophages before or during plating affected secretion of inflammatory molecules, like NO and IL-6, by these cells and if activation status of the macrophages had an effect on production of the adipocyte-specific inflammatory adipokine leptin. Results from these experiments can help clarify the role of macrophage-derived adipokines on the initiation of the macrophage-adipocyte inflammatory loop observed in obese adipose tissue.

We did not observe significant difference in NO or IL-6 production from macrophages regardless of if LPS was added before plating or during plating onto Transwells. As expected, increased NO production by the macrophages, a sign of macrophage activation, was accompanied by increased production of IL-6. Release of IL-6 by macrophages suggests that these macrophages are taking on an M1 phenotype, which is expected during LPS activation (31). Although the three methodologies of macrophage activation tested did not result in significant differences in NO and IL-6 production, it is important to note that at the time of cell-conditioned media collection, the media from macrophages plated in conditioned media (3T3+StimRAW+CondMed) contained secreted products for a 30 h time period while macrophages in fresh media (3T3+StimRAW) or with LPS added at time of plating (3T3+RAW+LPS) contained 24 h of secreted products (Figures 2 and 3). Levels of NO produced by activated macrophages plated in fresh media were similar to levels of NO produced by unstimulated macrophages suggesting that in this specific cell culture system these cells produced basal amounts of NO without LPS activation. This could be explained by activation of the macrophages as they are transferred from their original culture system (6-well plate) to the Transwells via the cell scraping method. On the other hand, quantified IL-6 levels were significantly higher in cell-conditioned media collected from activated macrophages plated in fresh media compared to unstimulated macrophages indicating that in this cell culture methodology challenging RAW 264.7 cells with LPS has a more pronounced and/or prolonged effect on production and release of IL-6 compared to NO.

In conclusion, the presence of LPS-stimulated macrophages in the co-incubation system did not affect leptin release by the mature adipocytes, as the adipocytes produced similar leptin levels when co-incubated with activated macrophages (3T3+RAW+LPS, 3T3+StimRAW, 3T3+StimRAW+CondMed) as when co-incubated with unstimulated macrophages (3T3+RAW) (Figure 4). Our results also highlight the importance of sampling and analyzing the macrophage and adipocyte-containing microenvironments (Transwells and wells, respectively) separately in order to detect the subtle signaling dynamics that are important in the paracrine conversation occurring between these cell types. The methodologies presented here can be adopted for the study of macrophage-adipocyte interactions, including cellular communication, chemotaxis studies, and effects of macrophage-derived molecules on adipocyte differentiation and mitochondrial function, among other research areas. Constant et al. (32) stated that the ERK 1/2-driven antiadipogenic effect of macrophage cell-conditioned media on adipocytes occurred during the first 2 days of the 8-day adipocyte differentiation period. The co-incubation protocols we have developed can be modified to test cellular communication between these two cell types at different time points, allowing for further analyses of temporal interactions. Transwell inserts with larger pore sizes (3-5 μm) can be used for migration and chemotaxis studies that can help answer questions about macrophages recruitment into obese AT, which appears to be one of the early steps in setting up the macrophage-adipocyte inflammatory loop.

## Acknowledgements

We are indebted to Dr. Andy McCall of Denison University’s Biology Department for his help with the statistical analyses.

## Author Contributions

Conceptualization: CCT

Data Curation: CCT & JB

Formal Analysis: CCT

Funding Acquisition: CCT & JB

Investigation: CCT & JB

Methodology: JB

Project Administration: CCT

Resources: CCT

Supervision: CCT

Visualization: CCT

Validation: CCT

Writing – original draft: CCT

Writing – review & editing: CCT

## Notes

### Competing Interest Statement

The authors have declared no competing interest.

## References

1. Ouchi N, Parker JL, Lugus JJ, Walsh K. Adipokines in inflammation and metabolic disease. Nat Rev Immunol. 2011 Feb;11(2):85–97. https://doi.org/10.1038/nri2921.

2. Sorisky A, Molgat ASD, Gagnon A. Macrophage-Induced Adipose Tissue Dysfunction and the Preadipocyte: Should I Stay (and Differentiate) or Should I Go?123. Adv Nutr. 2013 Jan 4;4(1):67–75.https://doi.org/10.3945/an.112.003020.

3. Fain JN. Release of Inflammatory Mediators by Human Adipose Tissue Is Enhanced in Obesity and Primarily by the Nonfat Cells: A Review. Mediators Inflamm [Internet]. 2010 [cited 2021 Jun 9];2010. https://doi.org/10.1155/2010/513948.

4. van Dielen F, van’t Veer C, Schols A, Soeters P, Buurman W, Greve J. Increased leptin concentrations correlate with increased concentrations of inflammatory markers in morbidly obese individuals. Int J Obes. 2001 Dec;25(12):1759–66. https://doi.org/10.1038/sj.ijo.0801825.

5. Friedman JM, Halaas JL. Leptin and the regulation of body weight in mammals. Nature. 1998 Oct;395(6704):763–70. https://doi.org/10.1038/27376.

6. Tartaglia LA, Dembski M, Weng X, Deng N, Culpepper J, Devos R, et al. Identification and expression cloning of a leptin receptor, OB-R. Cell. 1995 Dec 29;83(7):1263–71. https://doi.org/10.1016/0092-8674(95)90151-5.

7. Loffreda S, Yang SQ, Lin HZ, Karp CL, Brengman ML, Wang DJ, et al. Leptin regulates proinflammatory immune responses. The FASEB Journal. 1998;12(1):57–65. https://doi.org/10.1096/fsb2fasebj.12.1.57.

8. Mancuso P, Gottschalk A, Phare SM, Peters-Golden M, Lukacs NW, Huffnagle GB. Leptin-Deficient Mice Exhibit Impaired Host Defense in Gram-Negative Pneumonia. J Immunol. 2002 Apr 15;168(8):4018–24. https://doi.org/10.4049/jimmunol.168.8.4018.

9. Santos-Alvarez, J, Goberna, R, Sánchez-Margalet V. Human leptin stimulates proliferation and activation of human circulating monocytes. Cell Immunol. 1999 May 25;194(1):6–11. https://doi.org/10.1006/cimm.1999.1490.

10. Acedo SC, Gambero S, Cunha FGP, Lorand-Metze I, Gambero A. Participation of leptin in the determination of the macrophage phenotype: an additional role in adipocyte and macrophage crosstalk. In Vitro CellDevBiol-Animal. 2013 Jun;49(6):473–8. https://doi.org/10.1007/s11626-013-9629-x.

11. Unno Y, Akuta T, Sakamoto Y, Horiuchi S, Akaike T. Nitric oxide-induced downregulation of leptin production by 3T3-L1 adipocytes. Nitric Oxide. 2006 Sep 1;15(2):125–32. https://doi.org/10.1016/j.niox.2005.12.002.

12. Makki K, Froguel P, Wolowczuk I. Adipose Tissue in Obesity-Related Inflammation and Insulin Resistance: Cells, Cytokines, and Chemokines. ISRN Inflamm [Internet]. 2013 Dec 22 [cited 2021 Jun 9];2013. https://doi.org/10.1155/2013/139239.

13. Bai Y, Sun Q. Macrophage recruitment in obese adipose tissue. Obes Rev. 2015 Feb;16(2):127–36. https://doi.org/10.1111/obr.12242.

14. Osborn O, Olefsky JM. The cellular and signaling networks linking the immune system and metabolism in disease. Nat Med. 2012 Mar;18(3):363–74. https://doi.org/10.1038/nm.2627.

15. Boutens L, Stienstra R. Adipose tissue macrophages: going off track during obesity. Diabetologia. 2016;59:879–94. https://doi.org/10.1007/s00125-016-3904-9.

16. Lumeng CN, Bodzin JL, Saltiel AR. Obesity induces a phenotypic switch in adipose tissue macrophage polarization. J Clin Invest. 2007 Jan 2;117(1):175–84. https://doi.org/10.1172/JCI29881.

17. Yamashita A, Soga Y, Iwamoto Y, Asano T, Li Y, Abiko Y, et al. DNA microarray analyses of genes expressed differentially in 3T3-L1 adipocytes co-cultured with murine macrophage cell line RAW264.7 in the presence of the toll-like receptor 4 ligand bacterial endotoxin. Int J Obes. 2008 Nov;32(11):1725–9. https://doi.org/10.1038/ijo.2008.153.

18. Blüher M. The distinction of metabolically ‘healthy’ from ‘unhealthy’ obese individuals. Current Opinion in Lipidology. 2010 Feb;21(1):38–43. https://doi.org/10.1097/MOL.0b013e3283346ccc.

19. Arner E, Mejhert N, Kulyté A, Balwierz PJ, Pachkov M, Cormont M, et al. Adipose Tissue MicroRNAs as Regulators of CCL2 Production in Human Obesity. Diabetes. 2012 Aug 1;61(8):1986–93. https://doi.org/10.2337/db11-1508.

20. Kitade H, Sawamoto K, Nagashimada M, Inoue H, Yamamoto Y, Sai Y, et al. CCR5 Plays a Critical Role in Obesity-Induced Adipose Tissue Inflammation and Insulin Resistance by Regulating Both Macrophage Recruitment and M1/M2 Status. Diabetes. 2012 Jul 1;61(7):1680–90. https://doi.org/10.2337/db11-1506.

21. Patel P, Abate N. Body Fat Distribution and Insulin Resistance. Nutrients. 2013 Jun 5;5(6):2019–27. https://doi.org/10.3390/nu5062019.

22. Heilbronn LK, Rood J, Janderova L, Albu JB, Kelley DE, Ravussin E, et al. Relationship between Serum Resistin Concentrations and Insulin Resistance in Nonobese, Obese, and Obese Diabetic Subjects. The Journal of Clinical Endocrinology & Metabolism. 2004 Apr 1;89(4):1844–8. https://doi.org/10.1210/jc.2003-031410.

23. Vieira-Potter VJ. Inflammation and macrophage modulation in adipose tissues.Cellular Microbiology. 2014;16(10):1484–92. https://doi.org/10.1111/cmi.12336.

24. Zebisch K, Voigt V, Wabitsch M, Brandsch M. Protocol for effective differentiation of 3T3-L1 cells to adipocytes. Analytical Biochemistry. 2012 Jun 1;425(1):88–90. https://doi.org/10.1016/j.ab.2012.03.005.

25. Dobashi, K., et al. “Troglitazone Inhibits the Expression of Inducible Nitric Oxide Synthase in Adipocytes in Vitro and in Vivo Study in 3T3-L1 Cells and Otsuka Long-Evans Tokushima Fatty Rats.” Life Sciences, vol. 67, no. 17, Sept. 2000, pp. 2093–101. https://doi.org/10.1016/s0024-3205(00)00796-7.

26. Chirumbolo S, Franceschetti G, Zoico E, Bambace C, Cominacini L, Zamboni M. LPS response pattern of inflammatory adipokines in an in vitro 3T3-L1 murine adipocyte model. Inflamm Res. 2014 Jun 1;63(6):495–507. https://doi.org/10.1007/s00011-014-0721-9.

27. Jann B, Reske K, Jann K. Heterogeneity of Lipopolysaccharides. Analysis of Polysaccharide Chain Lengths by Sodium Dodecylsulfate-Polyacrylamide Gel Electrophoresis.European Journal of Biochemistry. 1975;60(1):239–46. https://doi.org/10.1111/j.1432-1033.1975.tb20996.x.

28. Yamashita A, Soga Y, Iwamoto Y, Yoshizawa S, Iwata H, Kokeguchi S, et al. Macrophage-Adipocyte Interaction: Marked Interleukin-6 Production by Lipopolysaccharide. Obesity. 2007;15(11):2549–52. https://doi.org/10.1038/oby.2007.305.

29. Vaughan T, Li L. Molecular mechanism underlying the inflammatory complication of leptin in macrophages. Molecular Immunology. 2010 Sep 1;47(15):2515–8. https://doi.org/10.1016/j.molimm.2010.06.006.

30. Olefsky JM, Glass CK. Macrophages, Inflammation, and Insulin Resistance.Annual Review of Physiology. 2010;72(1):219–46. https://doi.org/10.1146/annurev-physiol-021909-135846.

31. Orecchioni M, Ghosheh Y, Pramod AB, Ley K. Macrophage Polarization: Different Gene Signatures in M1(LPS+) vs. Classically and M2(LPS–) vs. Alternatively Activated Macrophages. Front Immunol. 2019 May 24 [cited 2021 Jun 9];10. https://doi.org/10.3389/fimmu.2019.01084.

32. Constant VA, Gagnon A, Yarmo M, Sorisky A. The antiadipogenic effect of macrophage-conditioned medium depends on ERK1/2 activation. Metabolism. 2008 Apr 1;57(4):465–72. http://dx.doi.org/10.1016/j.metabol.2007.11.005

